# FGF signaling promotes precursor spreading for adult adipogenesis in *Drosophila*

**DOI:** 10.1101/2022.04.21.489019

**Authors:** Yuting Lei, Yuwei Huang, Ke Yang, Xueya Cao, Yuzhao Song, Enrique Martín-Blanco, José C. Pastor-Pareja

**Affiliations:** School of Life Sciences, Tsinghua University, Beijing, 100084, China; Instituto de Biología Molecular de Barcelona, Consejo Superior de Investigaciones Científicas, Parc Científic de Barcelona, Barcelona, 08028, Spain; Tsinghua-Peking Center for Life Sciences, Beijing, 100084, China

**Keywords:** adipocyte, fat body, adipose tissue, migration, FGFR, lipodystrophy, autophagy, starvation

## Abstract

Knowledge of adipogenetic mechanisms is essential to understand and treat conditions affecting organismal metabolism and adipose tissue health. In *Drosophila*, mature adipose tissue (fat body) exists in larvae and adults. In contrast to the well-known development of the larval fat body from the embryonic mesoderm, adult adipogenesis has remained mysterious. Furthermore, conclusive proof of its physiological significance is lacking. Here, we show that the adult fat body originates from a pool of undifferentiated mesodermal precursors that migrate from the thorax into the abdomen during metamorphosis. Through in vivo imaging, we found that these precursors spread from the ventral midline and cover the inner surface of the abdomen in a process strikingly reminiscent of embryonic mesoderm migration, requiring FGF signaling as well. FGF signaling guides migration dorsally and regulates adhesion to the substrate. After spreading is complete, precursor differentiation involves fat accumulation and cell fusion that produces mature binucleate and tetranucleate adipocytes. Finally, we show that flies where adult adipogenesis is impaired by knock down of FGF receptor Heartless or transcription factor Serpent display ectopic fat accumulation in oenocytes and decreased resistance to starvation. Our results reveal that adult adipogenesis occurs de novo during metamorphosis and demonstrate its crucial physiological role.

## INTRODUCTION

Eukaryotic cells can efficiently store energy in the form of fat inside lipid droplets. Lipid droplets are ER outgrowths consisting of a core of neutral lipids surrounded by a phospholipid monolayer (Olzmann and Carvalho, 2019). Many unicellular eukaryotes and certain cell types in multicellular ones possess the ability to produce lipid droplets. However, in the animal kingdom, both vertebrates and arthropods have concentrated lipid storage and release functions in large specialized cells called adipocytes. Differentiated adipocytes associate into adipose tissues and display giant lipid droplets that occupy most of their cytoplasm. In vertebrates, histogenesis of adipose tissue (adipogenesis) is quite complex. Vertebrate adipocytes are generally believed to be of mesodermal origin, but specific populations have been found to derive instead from the neural crest (Le Lièvre and Le Douarin, 1975). In addition to fully differentiated adipocytes, mammalian adipose tissues contain stem cell precursors capable of producing new adipocytes (Berry et al., 2013; Spalding et al., 2008). Adipose tissue remodeling through formation of new adipocytes (hyperplasia) is a healthy response to caloric excess, whereas expansion of existing adipocytes through increased fat storage (hypertrophy) stresses those cells and associates with metabolic disease (Gupta, 2014). In contrast to adipocyte hyperplasia or hypertrophy, lipodystrophies are a group of congenital and acquired disorders characterized by the absence of functional adipocytes, causing insulin resistance, hyperlipidemia and other metabolic complications (Brown et al., 2016). Better knowledge of basic adipogenetic mechanisms, therefore, is essential to understand and treat conditions affecting adipose tissue health.

Besides vertebrates, the existence of true adipocytes and adipose tissue is documented in arthropods. The adipose tissue of arthropods, called fat body, has been extensively studied in insects, but is also present in crustaceans, chelicerates (spiders, scorpions and mites) and myriapods (Coons, 2014). Within insects, research in the fruit fly *Drosophila melanogaster* has described the presence of mature fat body in two stages of the life cycle of the animal: the larva and the adult. The development of the larval fat body, known to originate from the embryonic mesoderm, is well characterized. Shortly after gastrulation, at embryonic stage 11, two mesodermal cell layers become discernible: an internal one, in contact with the yolk sac, becomes the visceral musculature; an external one, lining the ectoderm, gives rise to the somatic musculature and the fat body (Campos-Ortega and Hartenstein, 1997). The transcription factor Srp is expressed in this embryonic fat body (Abel et al., 1993) and required for its development (Sam et al., 1996). During the larval stages, larval fat body adipocytes increase their cell size and ploidy through nutrition-dependent endoreplication (Britton and Edgar, 1998). Later, the larval fat body undergoes cell dissociation and histolysis during metamorphosis (Jia et al., 2014; Nelliot et al., 2006). Isolated larval fat body cells are found inside the adult abdomen until two days after eclosion. However, in addition to the disappearing larval adipocytes, the eclosed adult displays segmental plates of adult fat body lining the abdomen (Fig. 1A), with lesser amounts found in the head, thorax and female gonads. In contrast to the well-known development of the larval fat body, adult fat body adipogenesis has remained mysterious to this date (Hoshizaki et al., 1995; Li et al., 2019; Parra-Peralbo et al., 2021). Clonal analysis shows that the adult fat body, like the larval fat body, is mesodermal in origin (Lawrence and Johnston, 1986). A possible relation between the larval and adult adipose tissues has been a matter of speculation and discussion for long time. Two alternative mechanisms for adult adipogenesis have been proposed: partial reassociation of the dissociated larval fat body (Larsen, 1976) or de novo adipogenesis from undifferentiated precursors (Haunerland and Shirk, 1995). Many functional studies support a crucial involvement of the larval fat body in energy storage and metabolic regulation in the fast-feeding larva. Fewer studies, however, have tried to address the role of the adipose tissue in mature adult flies. Furthermore, due to insufficient knowledge of the development of the adult fat body and the lack of genetic tools to specifically image and manipulate it, conclusive proof of its physiological significance has been awaiting.

**Fig. 1.**
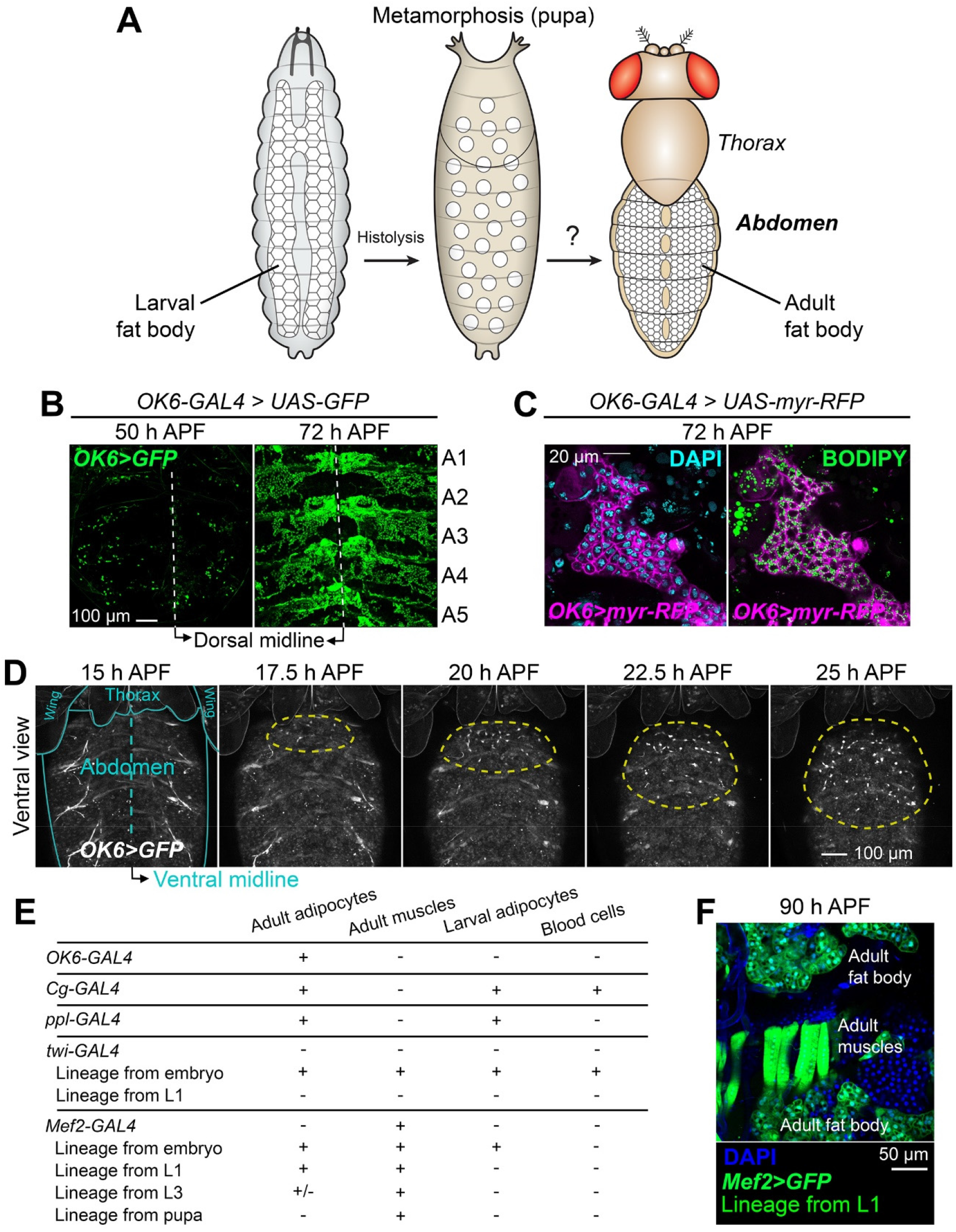
Adult fat body precursors migrate into the abdomen during metamorphosis. (A) Schematic cartoon depicting the adipose tissue (fat body) in *Drosophila* larvae and adults. The larval fat body, originating from embryonic mesoderm, disintegrates during metamorphosis. It is not known if the adult fat body is built from larval fat body remnants or assembled de novo from an alternate source of adipocytes. (B) Expression of GFP (green) driven by *OK6-GAL4* in pupal abdomens dissected and mounted flat at 50 and 72 h APF (after puparium formation). Abdominal segments A1 to A5 are indicated. (C) Adult fat body marked with *OK6-GAL4*-driven myr-RFP (magenta) at 72 h APF. Nuclei stained with DAPI (cyan, left) and neutral lipids with BODIPY (green, right). (D) Still images from a time-lapse recording of an *OK6>GFP* pupa (ventral view). To image the ventral abdomen, legs were gently displaced anteriorly. Blue lines outline wings, thorax and abdomen. Yellow dashed lines surround the growing population of *OK6*-positive cells migrating into the abdomen and proliferating there. Images are maximum intensity projections of 61 confocal sections. See Video S1. (E) Summary of GAL4 expression patterns, indicating the presence (+) or absence (-) of GFP expression under control of *OK6-GAL4, Cg-GAL4, ppl-GAL4, twi-GAL4* and *Mef2-GAL4* in different mesodermal derivatives. Shown as well are the results of lineage tracing experiments in which the progeny of all cells that express *twi-GAL4* or *Mef2-GAL4* at a given stage become permanently labeled (see Methods). (F) Abdomen of a pupa dissected 90 h APF in which cells that have expressed *Mef2-GAL4* up to the L1 stage are labeled with GFP (green). Summarized genotype: *Mef2-GAL4 + tub-GAL80*^*ts*^ *> UAS-Flp > act-y+-GAL4 > UAS-GFP*. Nuclei stained with DAPI (blue).

In this study, we set out to investigate the development of the adult fat body in *Drosophila*. Through in vivo imaging, we found that the adult fat body originates from a pool of precursors that migrate from the thorax into the abdomen during metamorphosis. These precursors spread from the ventral midline in a process strikingly reminiscent of embryonic mesoderm migration, requiring FGF signaling as well. In addition, we show that flies in which adult fat body development is impaired display decreased resistance to starvation.

## RESULTS

### Adult fat body precursors migrate into the abdomen during metamorphosis

Searching for tools that could help investigate the development of the adult fat body (Fig. 1A), we came across *OK6-GAL4*, a GAL4 enhancer trap insertion in the second chromosome (Sanyal, 2009). When we crossed *OK6-GAL4* to *UAS-GFP* flies, we observed expression of GFP in the pupal abdomen during metamorphosis. At 72 h APF (after puparium formation), *OK6*-driven GFP did not label the cells of the dissociated larval fat body, but was visible in segmental plates of tissue reminiscent of the morphology of the adult fat body (Fig. 1B). Staining with the neutral lipid dye BODIPY showed that *OK6*-positive cells contained lipid droplets, consistent with their identity as developing adipocytes (Fig. 1C). Because at earlier stages of metamorphosis *OK6*-driven GFP was expressed in single cells attached to the abdominal epidermis, we hypothesized that these were the precursors of the adult fat body. Time-lapse imaging of the pupa (see Methods) revealed that *OK6*-positive cells started arriving from the thorax at around 15 h APF, migrating and proliferating on the ventral epidermis of the abdomen (Fig. 1D; Video S1). To investigate the origin of these cells, we performed lineage tracing experiments. In these, GAL4-driven expression of the recombinase Flp, together with a flip-out cassette and thermosensitive GAL4 repressor GAL80^ts^, labeled the progeny of cells expressing at a given stage *twi-GAL4* (embryonic mesoderm) and *Mef2-GAL4* (myoblasts and muscles) (see Methods). Lineage tracing with *twi-GAL4* in the embryo labeled adult adipocytes (Fig. 1E), confirming their mesodermal origin (Lawrence and Johnston, 1986). Remarkably, *Mef2-GAL4* tracing in larva 1 (L1) stage labeled adult (but not larval) adipocytes and adult muscles (Fig. 1F). This result shows that adult and larval adipocyte lineages have diverged at the L1 stage and additionally suggests that adult adipocytes and muscles may descend from a common larval population of progenitors. Altogether, our data show that the adult fat body derives from a population of undifferentiated mesodermal precursors that migrate from the thorax into the abdomen during metamorphosis.

### The GATA transcription factor Serpent is required for early amplification of adult fat body precursors

Expression of the transcription factor Serpent (Srp) marks the precursors of the larval fat body in the embryo (Abel et al., 1993). Furthermore, *srp* mutant embryos lack fat body, indicating that Srp is essential for correct development of the larval fat body (Sam et al., 1996). We stained pupal abdomens with anti-Srp antibody and found that Srp was expressed in the adult fat body precursors, labeled with *OK6-GAL4*-driven GFP expression, and localized to their nuclei (Fig. 2A), hinting an involvement of Srp in adult fat body formation as well. In order to test this, we knocked down the expression of *srp* in the precursors of the adult fat body using *OK6-GAL4*-driven transgenic RNAi. In the abdomen of both wild type and *OK6>srp*^*i*^ adults dissected 2 days after eclosion, BODIPY staining revealed the presence of some fat body tissue. However, compared to wild type adults, the fat body of *OK6>srp*^*i*^ adults was severely reduced (Fig. 2B). To investigate the genesis of this phenotype, we imaged the adult fat body precursors using *OK6-GAL4*-driven GFP expression. We found that precursors were present in the ventral abdomen of *OK6>srp*^*i*^ pupae at 30 h APF (Fig. 2C). However, in contrast to the fast proliferation of these cells observed in the wild type, the number of precursors had increased less when we analyzed the same animal 6 h later at 36 h APF (Fig. 2D). From these results, we conclude that Srp expression in adult fat body precursors is necessary for the amplification of their numbers during the early phases of adult adipogenesis. Our results additionally suggest that Srp may not be involved in their correct specification and differentiation into adipocytes.

**Fig. 2.**
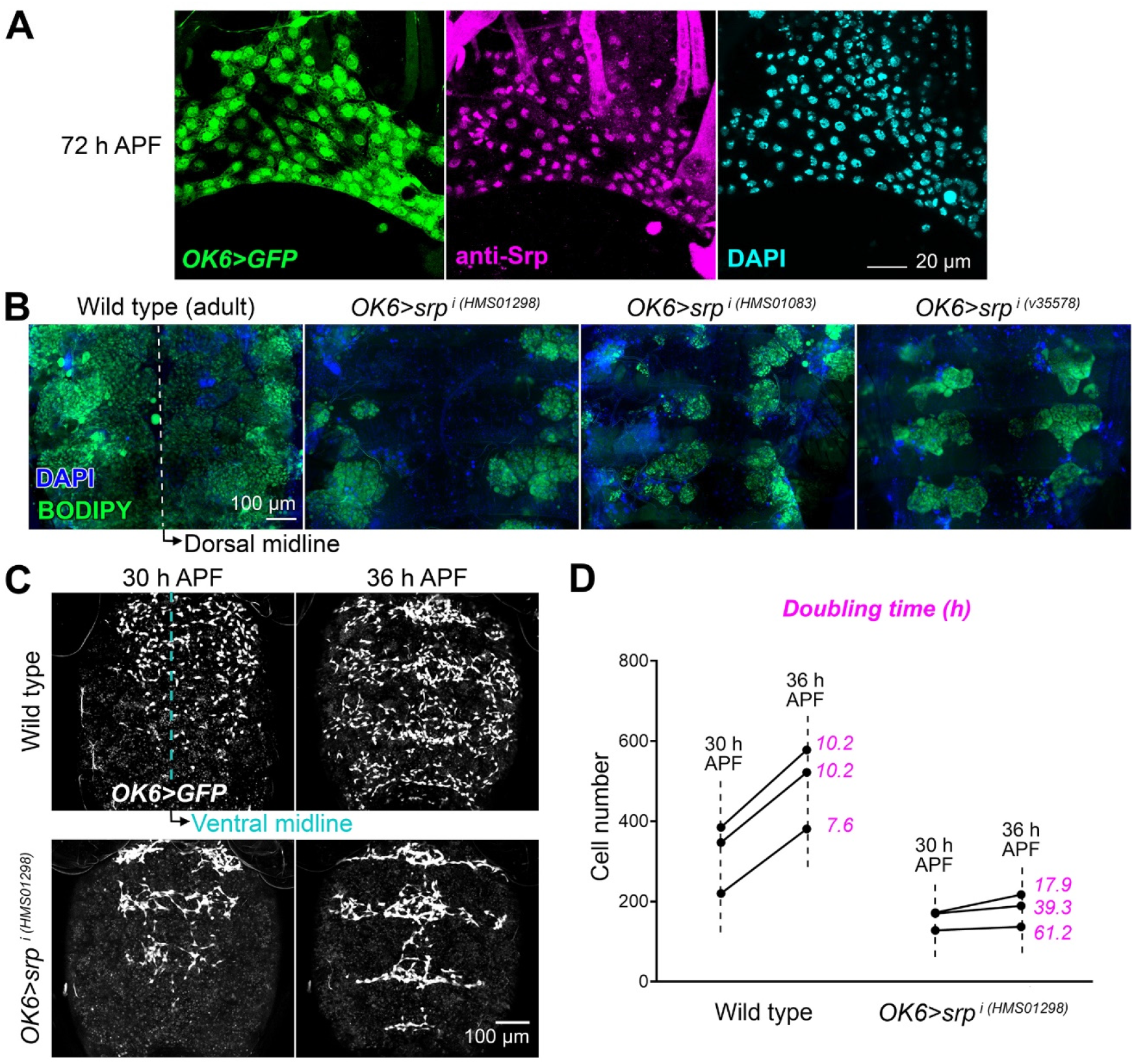
The GATA transcription factor Serpent is required for amplification of adult fat body precursors. (A) Adult fat body precursors (*OK6-GAL4*-driven GFP, green) from an abdomen dissected 72 h APF and stained with anti-Srp antibody (magenta). Nuclei stained with DAPI (cyan). (B) Adult abdomens from a wild type fly and flies in which *srp* was knocked down with three different RNAi transgenes under *OK6-GAL4* control (O*K6>srp*^*i*^). Abdomens were dissected 2 days after eclosion and mounted flat after staining with DAPI (nuclei, blue) and BODIPY (neutral lipids, green). (C) Adult fat body precursors (*OK6-GAL4*-driven GFP, white) imaged in vivo in the abdomen (ventral view) of wild type (top) and O*K6>srp*^*i*^ (bottom) pupae at 30 (left) and 36 (right) h APF. Images are maximum intensity projections of 65 confocal sections. (D) Graph representing number of adult fat body precursors at 30 and 36 h APF in three wild type and three O*K6>srp*^*i*^ animals. Cells were counted in images like those in (C).

### Adult fat body precursors spread from the ventral midline

To further investigate adult adipogenesis, we recorded and analyzed the behavior of adult fat body precursors after the initial migration that brings them to the abdomen. Time lapse imaging of *OK6>GFP* animals from 30 h APF showed that precursors in the ventral abdomen quickly converge towards the ventral midline at about 32 h APF (Fig. 3A; Video S2). This convergent movement is coincident with the time when the expanding nests of histoblasts (adult epidermal cells) push and replace the contracting ventral epidermis of the larva (Ninov and Martín-Blanco, 2007). After this ventral contraction, adult fat body precursors spread on the abdominal epidermis, first laterally (Fig. 3A; Video S2) and then dorsally (Fig. 3B; Video S2), as they continue to increase their numbers through cell proliferation. Once the spreading precursors reach the dorsal side, they converge towards the dorsal midline from left and right (Fig. 3C; Video S2). Tracking of cell trajectories showed that adult fat body precursors experienced frequent changes of direction and repulsive interactions during their spreading when they contacted or collided with each other (Fig. 3D; Video S2). At the same time, analysis of the direction of migration with a 4-minute resolution (the recording interval of our movies) revealed a tendency towards displacements in the dorsal direction (Fig. 3E). Consistent with both contact inhibition and guided migration governing precursor movement, the observed long-term displacement of precursors is less dorsally oriented than predicted from the observed 4-min directional bias (Fig. 3E). In all, our time-lapse recordings show that adult fat body precursors spread throughout the abdomen from the ventral midline (Fig. 3F). In addition, our analysis of their trajectories indicates that a directional component guiding migration dorsally is operative besides mutual repulsion.

**Fig. 3.**
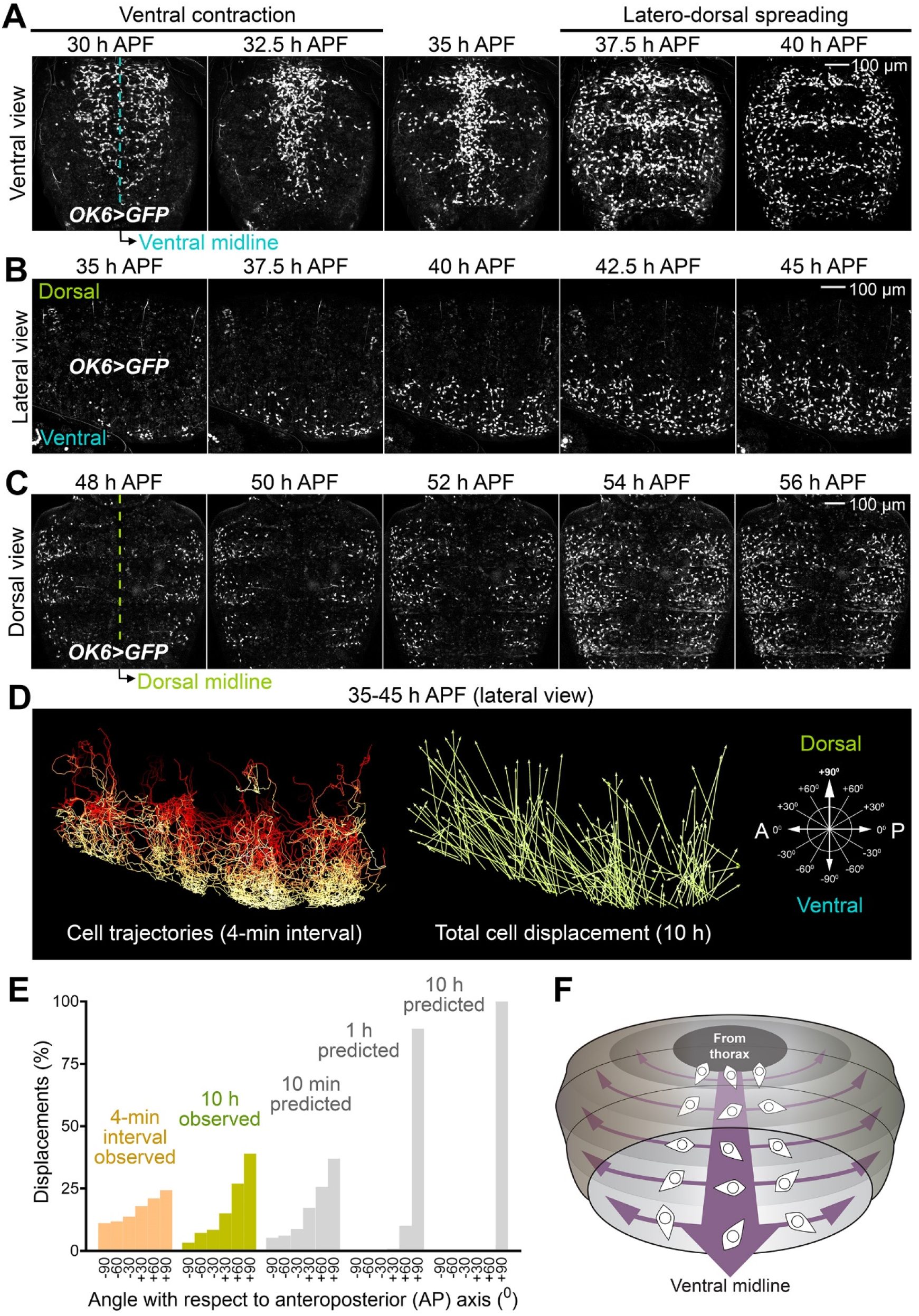
Adult fat body precursors spread from the ventral midline. (A) Still images from a time-lapse recording of adult fat body precursors (*OK6>*GFP, white) in the ventral abdomen of the pupa (30-40 h APF). Images are standard deviation projections of 62 confocal sections. See Video S2. (B) Still images from a time-lapse recording of adult fat body precursors (*OK6>*GFP, white) imaged laterally in the abdomen of the pupa (35-45 h APF; dorsal up, anterior left). Images are maximum intensity projections of 62 confocal sections. See Video S2. (C) Still images from a time-lapse recording of adult fat body precursors (*OK6>*GFP, white) in the dorsal abdomen of the pupa (48-56 h APF). Images are maximum intensity projections of 62 confocal sections. See Video S2. (D) Trajectories of the adult fat body precursors imaged in (B). On the left, complete migration paths are represented (a stack of images for each time point was acquired every 4 min). Total precursor displacement for the 10 h duration of recording is represented as well. See Video S2. (E) Graph representing the angle of migration of precursors in (B) with respect to the anterior-posterior axis. The angle of a fully dorsal displacement is 90° [see schematic representation in (D)]. For the 4-min interval angle distribution, n=22,831. For the 10 h angle distribution, n=485. Predicted distributions based on the assumption that migration is governed solely by dorsal displacement bias (no repulsion) were calculated by iteration of the observed 4-min interval angle distribution. (F) Schematic illustration of the migration of adult fat body precursors in the abdomen during metamorphosis. Precursors spread laterally and dorsally from the ventral midline.

### FGF signaling is required for adult adipogenesis

The spreading of adult fat body precursors from the ventral midline during metamorphosis, our live imaging showed, is very reminiscent of the migration of embryonic mesodermal cells during gastrulation, taking place after ventral furrow ingression (Leptin and Grunewald, 1990). Mesoderm migration in the embryo depends on Fibroblast Growth Factor (FGF) signaling. We therefore decided to test the involvement of FGF signaling in the adult fat body formation as well. Embryonic mesoderm cells express the FGF receptor Heartless (Htl) (Beiman et al., 1996; Gisselbrecht et al., 1996), whereas the overlying ectoderm expresses its FGF ligands Pyramus (Pyr) and Thisbe (Ths) (Gryzik and Müller, 2004; Stathopoulos et al., 2004) (Fig. 4A). Knock down of *htl* under control of *OK6-GAL4* in adult fat body precursors produced adults lacking most fat body tissue in their abdomens (Fig. 4B and C). Expression of a dominant negative version of Htl (Htl^DN^) similarly caused a large reduction in adult fat body tissue (Fig. 4D). Expression of a constitutively active Htl (Htl^CA^), in contrast, did not cause any apparent defect (Fig. 4E). Consistent with a requirement of *htl* in the formation of the adult fat body, a *htl-GAL4* reporter showed expression in the adult fat body precursors (Fig. 4F; Video S3). We next knocked down the expression of Htl-binding ligands Ths and Pyr under control of strong, ubiquitous driver *act-GAL4*. Knock down of *ths* strongly reduced the amount of fat body tissue in the adult abdomen (Fig. 4G). Knock down of *pyr*, in contrast, did not show such effect (Fig. 4H). Consistent with a requirement of *ths* in the formation of the adult fat body, a *ths-GAL4* reporter was expressed in the dorsal epidermis of the pupal abdomen (Fig. 4I). In all, these data evidence that expression of FGF receptor Htl in adult fat body precursors and FGF ligand Ths in the epidermis are needed for formation of the adult fat body.

**Fig. 4.**
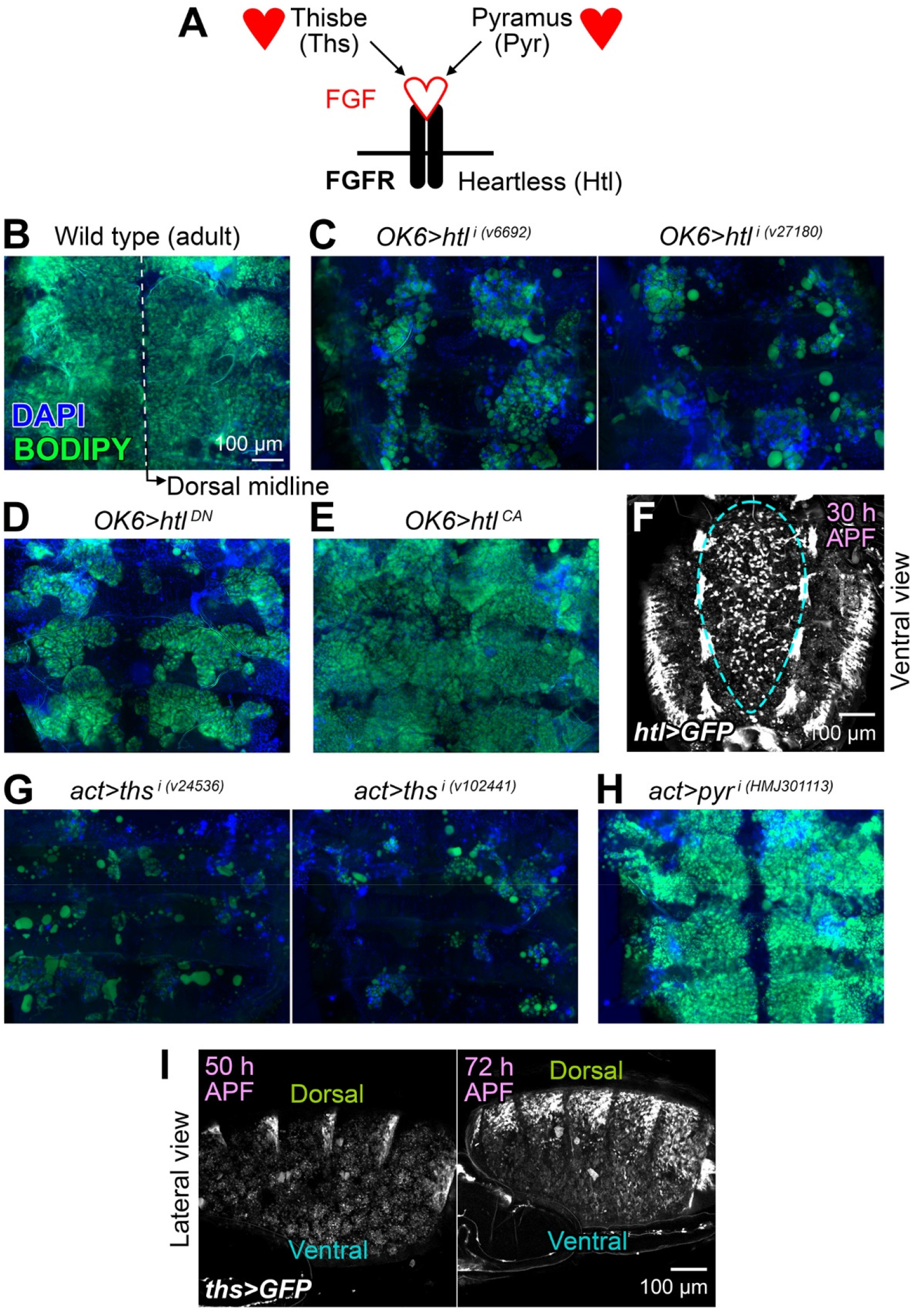
FGF signaling is required for adult adipogenesis. (A) Cartoon representing the receptor Heartless (htl) and its two known FGF-like ligands Thisbe (Ths) and Pyramus (Pyr). (B) Adult abdomens from a wild type fly dissected 2 days after eclosion and mounted flat after staining with DAPI (nuclei, blue) and BODIPY (neutral lipids, green). (C) Adult abdomens from flies in which *htl* was knocked down under control of *OK6-GAL4* (O*K6>htl*^*i*^) using two different RNAi transgenes. DAPI (blue) and BODIPY (green) stainings are shown. (D) Adult abdomen from a fly expressing dominant negative Htl (O*K6>htl*^*DN*^). DAPI (blue) and BODIPY (green) stainings are shown. (E) Adult abdomen from a fly expressing a constitutively active Htl (O*K6>htl*^*CA*^). DAPI (blue) and BODIPY (green) stainings are shown. (F) Expression of GFP (white) under control of *htl-GAL4* in motile fat body precursors imaged in vivo 30 h APF in the abdomen (ventral view). Images are maximum intensity projections of 60 confocal sections. See Video S3. (G) Adult abdomens from flies in which *ths* was knocked down under control of *act-GAL4* (*act>htl*^*i*^) using two different RNAi transgenes. DAPI (blue) and BODIPY (green) stainings are shown. (H) Adult abdomen from a fly in which *pyr* was knocked down under control of *act-GAL4* (*act>pyr*^*i*^). DAPI (blue) and BODIPY (green) stainings are shown. (I) Expression of GFP (white) under control of *ths-GAL4* in the dorsal epidermis of the abdomen (lateral view) at 50 and 72 h APF. Images are maximum intensity projections of 60 confocal sections.

### FGF signaling confers directionality and substrate adherence during precursor spreading

We next tried to ascertain the role of FGF signaling in adult adipogenesis. To that end, we imaged and analyzed the migration of adult fat body precursors in conditions of loss of FGF signaling (expressing Htl^DN^, Fig. 5A) and excess FGF signaling (expressing Htl^CA^, Fig. 5B). In both cases, live imaging showed precursors spreading (Fig. 5A-C; Video S4). Analysis of their direction of migration (4-min interval), however, revealed that expression of both Htl^DN^ and Htl^CA^ markedly reduced their tendency to displace dorsally (Fig. 5D). This result is highly consistent with a role of dorsally-expressed Ths in guiding the migration of Htl-expressing precursors as a chemoattractant cue. However, as previously noted, expression of Htl^CA^ in precursors produced a seemingly normal adult fat body (see Fig. 4E), unlike Htl^DN^ (Fig. 4D), hinting FGF roles additional or alternative to chemoatraction that might explain this discrepancy. To solve this, we further analyzed the spreading of *OK6>htl*^*DN*^ and *OK6>htl*^*CA*^ adult fat body precursors, and found that expression of Htl^CA^, but not Htl^DN^, reduced their migration speed (Fig. 5E). Furthermore, counting of precursors in lateral view recordings from 48 to 58 h APF revealed that the number of precursors expressing Htl^DN^ decreased over time (Fig. 5F), instead of increasing as a result of proliferation and income of new cells from the ventral side. Upon close observation, we discovered that precursors expressing Htl^DN^ frequently detached from the abdominal epidermis and abandoned the plane of view to disappear into the body cavity (Fig. 5G; Video S5). Altogether, our analysis of *htl* loss and gain of function phenotypes is consistent with a function of FGF signaling in both directing migration dorsally and increasing adhesion to the substrate.

**Fig. 5.**
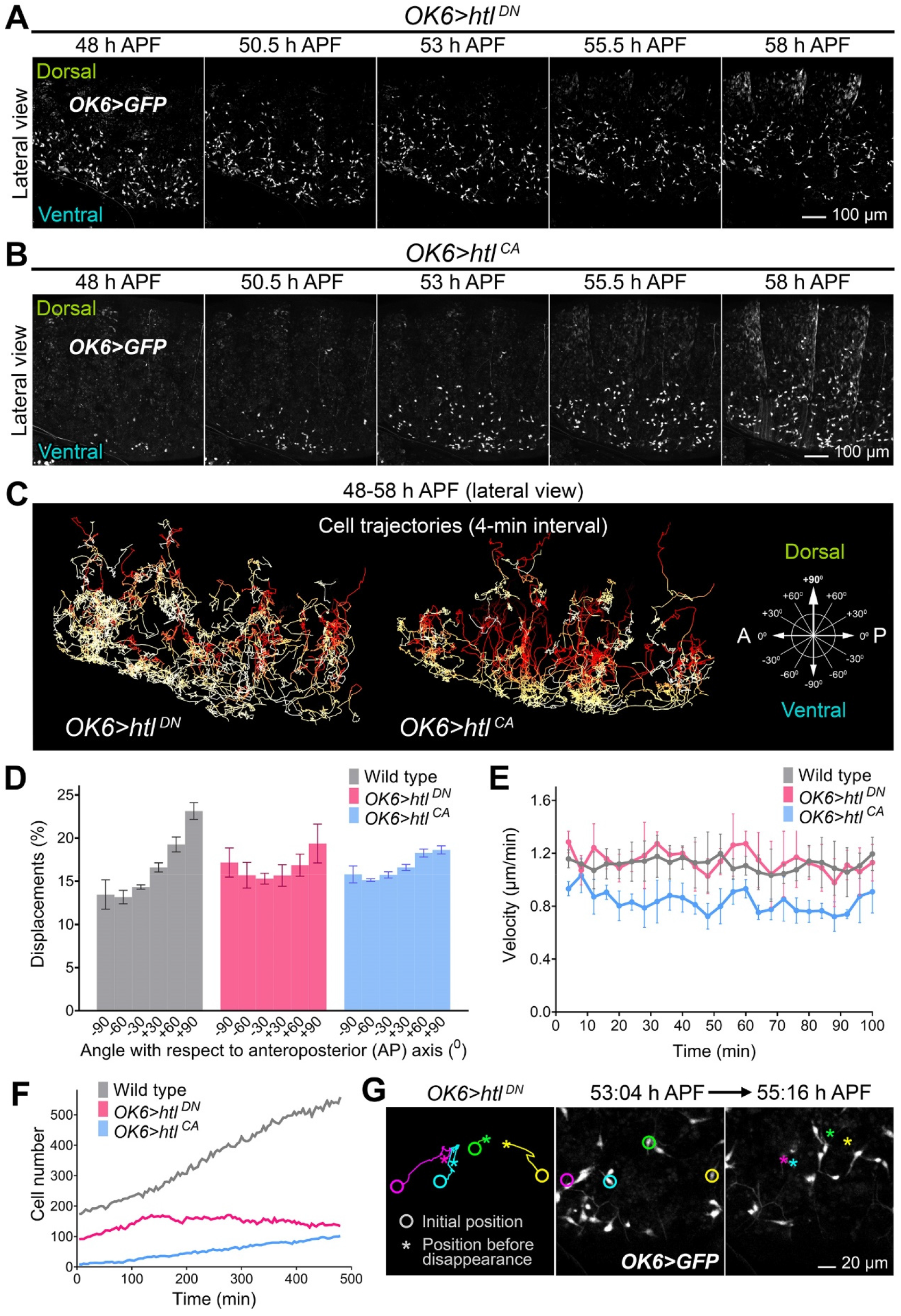
FGF signaling confers directionality and substrate adherence during precursor spreading. (A) Still images from a time-lapse recording of *OK6>htl*^*DN*^ adult fat body precursors (*OK6>GFP*, white) imaged laterally in the abdomen of the pupa (48-58 h APF; dorsal up, anterior left). Images are maximum intensity projections of 60 confocal sections. See Video S4. (B) Still images from a time-lapse recording of *OK6>htl*^*CA*^ adult fat body precursors (*OK6>GFP*, white) imaged laterally in the abdomen of the pupa (48-58 h APF; dorsal up, anterior left). Images are maximum intensity projections of 60 confocal sections. See Video S4. (C) Trajectories of the adult fat body precursors imaged in (A) and (B). Complete migration paths are represented (a stack of images for each time point was acquired every 4 min). See Video S4. (D) Angle of migration (4-min interval) with respect to the anterior-posterior axis of wild type, *OK6>htl*^*DN*^ and *OK6>htl*^*CA*^ precursors imaged laterally 52-54 h APF. Three recordings were analyzed for each genotype. Error bars represent SD. (E) Average migration velocity (4-min interval) of wild type, *OK6>htl*^*DN*^ and *OK6>htl*^*CA*^ precursors imaged laterally 52-54 h APF. Three recordings were analyzed for each genotype. Error bars represent SD. (F) Evolution of precursor numbers with time in wild type, *OK6>htl*^*DN*^ and *OK6>htl*^*CA*^ abdomens imaged laterally 48-58 h APF. Notice stationary/decreasing number of *OK6>htl*^*DN*^ precursors. (G) Still images from a time-lapse recording of *OK6>htl*^*DN*^ adult fat body precursors (*OK6>GFP*, white) imaged laterally in the abdomen of the pupa (dorsal up, anterior left) at 53:04 (center panel) and 55:16 h APF. Initial position, trajectory and final position before detachment of four precursors are represented in the left panel. Images are maximum intensity projections of 60 confocal sections. See Video S5.

### Adult fat body adipocytes are formed by fusion of precursors after spreading

As a result of spreading and continued proliferation, adult fat body precursors end up covering most of the inner surface of the abdominal epidermis, stop migrating and become confluent at about 65 h APF, giving rise to a tissue monolayer (Fig. 6A). Soon after becoming confluent, precursors start accumulating fat in the form of lipid droplets, suggesting a process of gradual differentiation into mature adipocytes, complete at day 2 after eclosion of the adult (Fig. 6B). During this differentiation process, we noticed the progressive appearance of adipocytes containing two and four nuclei (Fig. 6B), consistent with the observation of binucleate and tetranucleate adipocytes in the adult by others (Doane, 1960). Indeed, our counts showed that the adult fat body consists entirely of binucleate and tetranucleate cells in proportions that do not seem to change after eclosion (Fig. 6C). To ascertain the mechanism by which adult adipocytes become multinucleate, we imaged their late metamorphic development and documented multiple instances of cells merging through disappearance of the intervening plasma membranes (Fig. 6D; Video S6). These observations indicate that adult adipocytes become multinucleate not through mitosis followed by incomplete cytokinesis, as is the case in the mammalian liver (Guidotti et al., 2003), but as a result of cell fusion. In these binucleate and tetranucleate adipocytes, in addition, we found that the DNA content of nuclei in adults was 4C in average (Fig. 6E). Furthermore, adult adipocyte nuclei were surrounded by a cortex of perinuclear microtubules (Fig. 6F), typical of polyploid cells (Sun et al., 2019). This suggests that adult adipocyte nuclei have switched to an endoreplicative cell cycle and, therefore, are tetraploid rather than diploid stalled in G2. Our data show that adult fat body precursors give rise to large binucleate and tetranucleate adipocytes during late metamorphosis through cell fusion (Fig. 6G).

**Fig. 6.**
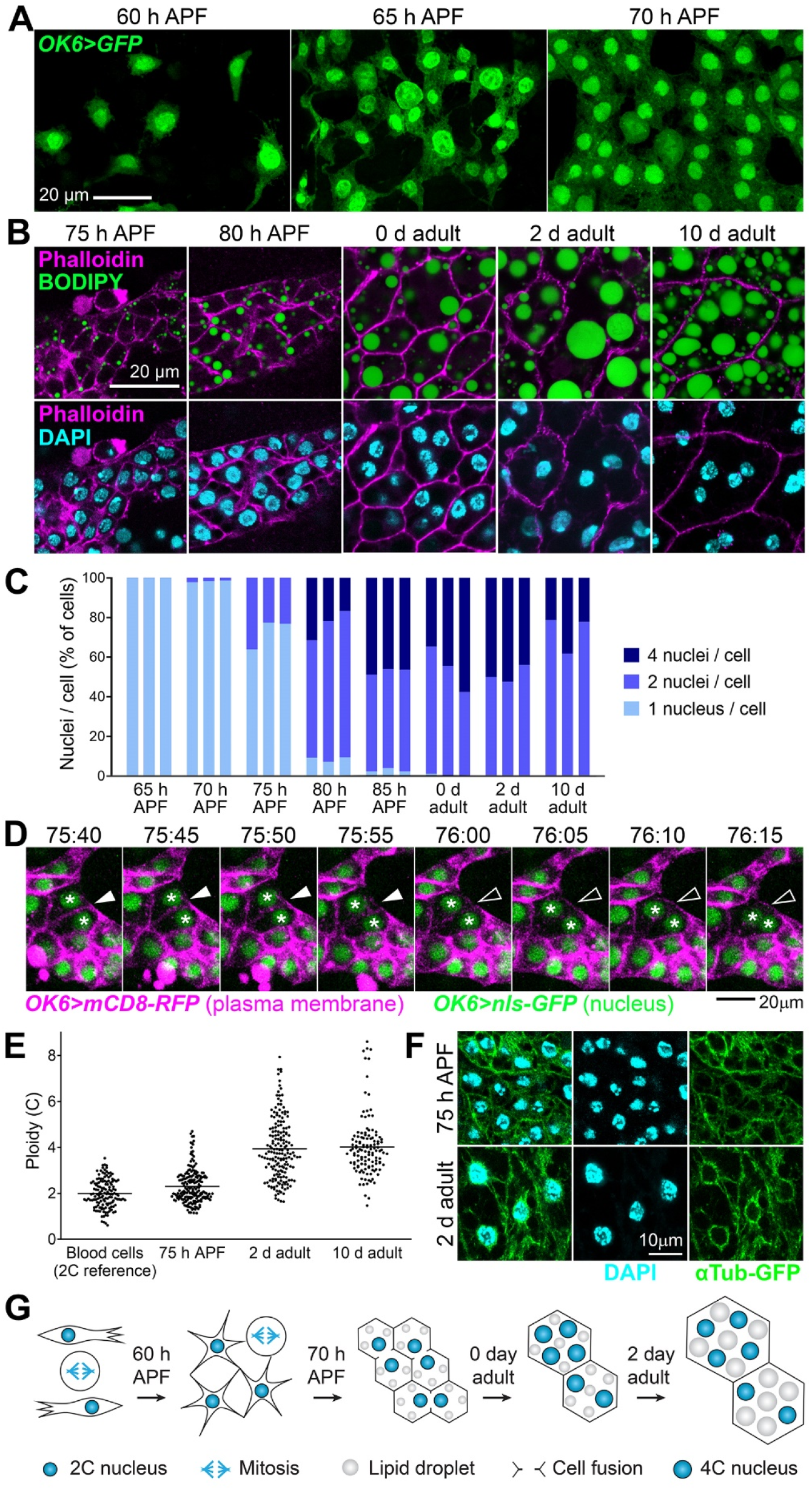
Adult fat body adipocytes are formed by fusion of precursors after spreading. (A) Adult fat body precursors (*OK6>GFP*, green) imaged at indicated times. Notice the presence of dividing cells. (B) Images showing the differentiation of adult precursors into mature adipocytes at indicated times. Eclosion of the adult takes place at about 96 h APF. Stainings with phalloidin (actin cell cortex, magenta, upper and lower row), BODIPY (neutral lipids, green, upper row) and DAPI (nuclei, cyan, lower row) are shown. Notice gradual fat accumulation, cell size increase and appearance of binucleate and tetranucleate cells until day 2 of eclosion. (C) Proportion of cells containing 1, 2 and 4 nuclei at indicated times. Counts in three individuals per time point are represented. At least 100 cells were analyzed per individual at 70 and 75 h APF, 40 cells at 80 and 85 h APF, and 30 cells in adults. (D) Still images from a time-lapse recording of the fusion of two adult fat body precursors at indicated times (75:40-76:15 h APF). Plasma membrane is labeled with mCD8-RFP (magenta) and nuclei with nls-GFP (green), both driven by *OK6-GAL4*. Arrowheads point to the disappearing membrane separating the two cells. Asterisks mark their nuclei. See Video S6. (E) Ploidy in nuclei of adult adipocytes at indicated times. Ploidy was estimated from confocal stacks by measuring the amount of DAPI signal through the entire nuclear volume (see Methods). Blood cells were used as a diploid reference (2n, 2C). Each point represents a measurement in a single nucleus. Horizontal lines mark the average value. (F) Microtubule organization in adult adipocytes at 75 h APF (upper row) and 2 days after eclosion (lower row). Microtubules are marked with αTub-GFP (green) driven by *OK6-GAL4* and *Cg-GAL4*, respectively. Nuclei stained with DAPI (cyan). Notice perinuclear microtubule organization in 2 day adult nuclei. (G) Cartoon depicting the maturation of adult fat body precursors into mature adipocytes.

### The adult fat body buffers fat levels and provides resistance to starvation

After studying the morphogenesis of the adult fat body during metamorphosis, we sought to get insights into its function and evidence of its physiological importance. To this end, we analyzed adult flies in which the adult fat body was missing or severely reduced due to knock down of *srp* or *htl* under *OK6-GAL4* control. In these flies, we found that neutral lipids ectopically accumulated in oenocytes (Fig. 7A), a cell type involved like the fat body in lipid metabolism (Makki et al., 2014). This result suggests a central role for the adult fat body in storing away fat and buffering its circulation levels in the animal. Further proof of an essential storage role for the adult fat body, its fat content decreased when we subjected adults to starvation for 3 days and recovered normal levels when flies thus starved were refed for 1 day (Fig. 7B). Reduction of fat levels upon starvation was accompanied by autophagy, as evidenced by the presence of vesicles positive for autophagy marker Atg8, reversible upon refeeding as well (Fig. 7C). These results strongly argue that the adult fat body acts as an energy reserve to be mobilized upon starvation. To finally probe the importance of this reserve, we recorded the survival after starvation of wild type flies and flies in which adult adipogenesis was impaired due to *srp* or *htl* knock down. Compared to control flies, survival of flies in which *srp* or *htl* had been knocked under *OK6-GAL4* control was reduced in approximately one day (Fig. 7D), showing that the adult fat body reserve provides increased resistance to starvation. In summary, we conclude that the adult fat body, formed de novo during metamorphosis, accomplishes a fat storage role crucial in the physiology of the adult.

**Fig. 7.**
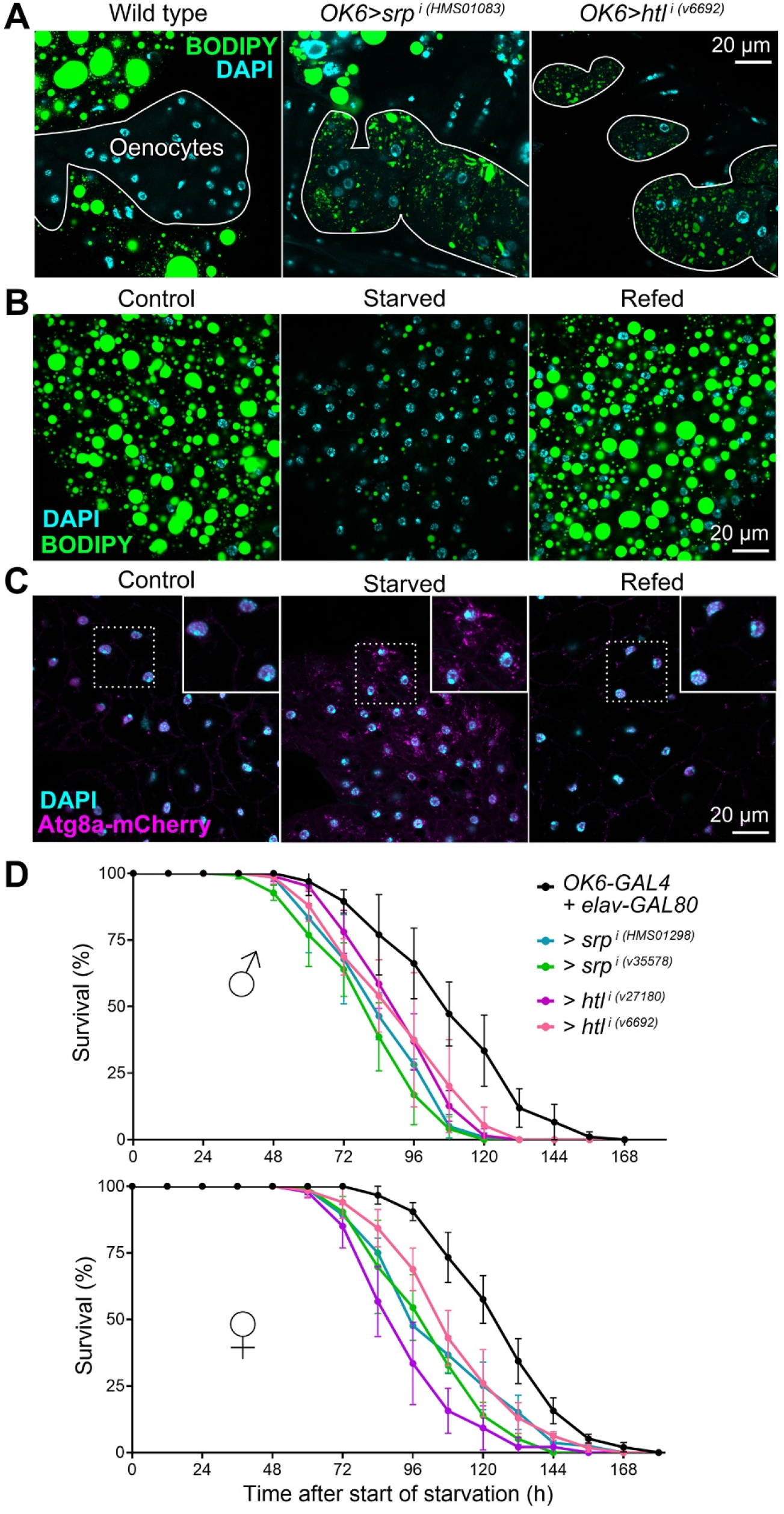
The adult fat body buffers fat levels and provides resistance to starvation. (A) Oenocytes (outlined) from adult wild type (left panel), *OK6>srp*^*i*^ (center panel) and *OK6>htl*^*i*^ (right panel) abdomens dissected 2 days after eclosion, stained with BODIPY (green) and DAPI (cyan). Notice the accumulation of lipid droplets in *OK6>srp*^*i*^ and *OK6>htl*^*i*^ oenocytes. (B) Adult fat body stained with BODIPY (green) in control flies (5 day adult, left panel), starved flies (2 day adult starved for 3 days, center panel) and refed flies (2 day adult starved for 3 days and refed for 1 day, right panel). Nuclei stained with DAPI (cyan) (C) Adult fat body expressing autophagy marker Atg8a-mCherry (driven by *BM-40-SPARC-GAL4*, magenta) in control flies (5 day adult, left panel), starved flies (2 day adult starved for 3 days, center panel) and refed flies (2 day adult starved for 3 days and refed for 1 day, right panel). Areas inside dashed squares are magnified in upper right corner insets. Nuclei stained with DAPI (cyan). (D) Graphs representing survival of control, *OK6>srp*^*i*^ and *OK6>htl*^*i*^ adult male (upper graph) and female (lower graph) flies subjected to starvation starting to 2 days after eclosion. *elav-GAL80*, repressing GAL4 activity in neurons, was included in genotypes to prevent a possible influence of *OK6-GAL4* expression in larval motoneurons (Sanyal, 2009). Three repeats were carried out per genotype and sex, each with at least 85 flies. Error bars represent SD. In all cases, differences with the control were significant in Mantel-Cox tests (****; p< 0.0001).

## DISCUSSION

In this study, we found that adult *Drosophila* adipocytes do not originate in the larval adipose tissue. Instead, de novo adipogenesis takes place during metamorphosis, when the adult fat body assembles from undifferentiated mesodermal precursors. Through in vivo imaging, we observed that fast-proliferating adult fat body precursors migrate from the thorax into the abdomen, accumulate at the abdominal ventral midline and spread laterally and dorsally on the inner surface of the abdominal epidermis. The migration of these precursors is, therefore, strikingly reminiscent of mesoderm migration during gastrulation in the embryo (Leptin and Grunewald, 1990; McMahon et al., 2008; Sun et al., 2020). During gastrulation, the cells that give rise to the mesoderm invaginate at the ventral midline, undergo epithelial-to-mesenchymal transition and migrate dorsally along the ectoderm. Outside of insects, a form of secondary gastrulation has been reported during metamorphosis of the jellyfish *Aurelia* (Yuan et al., 2008). Another metamorphic process in *Drosophila* with a clear counterpart in embryonic development is thorax closure at the dorsal midline, reminiscent of embryonic dorsal closure and driven as well by JNK activity in an epidermal edge (Martín-Blanco et al., 1998; Pastor-Pareja et al., 2004). The extent to which metamorphosis recapitulates key morphogenetic processes of embryonic developmentt is worth exploring. Indeed, comparisons among different *Drosophila* species revealed reduced transcriptome divergence during both mid-embryogenesis (Kalinka et al., 2010) and metamorphosis (Artieri and Singh, 2010), suggesting intense developmental constrains shared by both stages.

Supporting the notion that adult adipogenesis recapitulates embryonic mesoderm formation, we found that FGF signaling, required for mesoderm migration, is critical also for adult fat body formation. In both processes, epidermal FGF ligands activate FGF receptor Htl in motile precursors. In the embryo, FGF signaling mutants fail to spread their mesoderm from the ventral midline. Different roles have been attributed to FGF to explain this defect, such as promoting epithelial-to-mesenchymal transition, regulating proliferation and guiding migration as a chemoattractant cue (Muha and Müller, 2013; Sun et al., 2020). According to our results, FGF may not affect precursor proliferation or differentiation during adult adipogenesis. However, our findings show that FGF acts as a chemoattractant cue that influences the direction of migration, since both loss and gain of *htl* function makes precursor displacements less dorsally directed. In good agreement with such a guidance role, FGF Ths is expressed in the dorsal epidermis. Nonetheless, our analysis reveals that a second effect of FGF signaling, potentially more important for adult adipogenesis, is to enhance adhesion between precursors and the epidermis on which they migrate. This conclusion is based on (1) the behavior of *htl*^*DN*^ precursors, which frequently detach and disappear into the body cavity, (2) the decreased migration speed of *htl*^*CA*^ precursors, and (3) the fact that *htl*^*CA*^ precursors give rise to an apparently normal adult fat body despite decreased motility and directionality. In this light, mutual repulsion and contact inhibition of locomotion, similar to the dispersion of embryonic blood cells (Davis et al., 2012), may be sufficient to ensure spreading of *htl*^*CA*^ precursors, although at a slower pace. In embryonic mesoderm spreading, one of the defects reported in h*tl* mutants is reduced expression of *mys*, encoding a β subunit of the extracellular matrix receptor integrin (Sun and Stathopoulos, 2018). Reduced integrin adhesion could explain the detachment of Htl^DN^ precursors, but does not fit well their normal velocity, as loss of integrin should reduce migration speed. A recently described process in which FGF may predominantly regulate adhesion instead of acting as a guidance cue is the wrapping of olfactory glomeruli expressing Ths by Htl-expressing ensheathing glia (Wu et al., 2017). In these and other processes, the mechanisms by which FGF signaling regulates substrate adherence should be further investigated.

Another point for future clarification is the exact origin of adult fat body precursors and their location prior to migration into the abdomen. Lineage tracing experiments with *Mef2-GAL4* suggest a common lineage with the myoblast precursors of adult muscles. It has been proposed that adult fat body cells derive from adepithelial cells (Hoshizaki et al., 1995). These are populations of cells in the larval imaginal discs that contain large numbers of adult muscle precursors. According to the images in the reference, however, the putative precursor cells are not adepithelial cells, but differentiated blood cells that typically attach to other regions of the imaginal discs such as the wing pleura or the antenna. Despite this, we consider imaginal disc-associated adepithelial cells indeed a likely source of the adult adipocyte precursors. Similar to the adipogenic precursors, adepithelial myoblasts express Htl and respond to the expression of FGF ligands in the epidermis (Dutta et al., 2005; Everetts et al., 2021; Vishal et al., 2020). Further suggesting similarity between fat body and muscle precursors is production of syncitia through cell fusion. Adipocyte fusion, however, seems homotypical, unlike myoblast fusion, in which a founder asymmetrically instructs other cells to fuse with it (Lee and Chen, 2019). It will be interesting to investigate in the future how the temporal window of adipocyte cell fusion is determined (70 to 96 h APF) and whether the conserved machinery by which myoblasts fuse to produce muscle fibers is acting in adipocyte fusion as well.

The consequences and potential advantages for the function of adipocytes of their status as binucleate and tetranucleate polyploid cells is an additional topic of interest stemming from our findings. Human and rodent hepatocytes are frequently binucleate, but this is due to abortive cytokinesis rather than cell fusion (Guidotti et al., 2003). Moreover, in the human liver endoreplication produces tetraploid and octaploid nuclei, giving rise to a mixture of mononucleate 2n, 4n, 8n and binucleate 2×2n, 2×4n cells (Toyoda et al., 2005). Faster attainment of large cell volumes is often adduced to explain polyploidy in the larval fat body and other endoreplicating larval tissues (Orr-Weaver, 2015), a purpose cell fusion could serve as well. Alternatively, it has been proposed that polyploidy confers protection to human hepatocytes against genotoxic damage. Under this lens, acquiring multiple genome copies could buffer the effects of mutations caused by DNA damaging agents (Pandit et al., 2013). Regardless of the reasons behind multinucleation/polyploidy in the adult fat body, a possible consequence, our results suggest, is limited tissue plasticity. We did not observe after eclosion mitotic cells, nor any variation in nuclei number, DNA content (4C) or proportions of binucleate/tetranucleate cells. Starvation did not seem to induce changes in these features either. Only size increase was apparent from 0 to 2 days after eclosion, consistent with previous reports (Johnson and Butterworth, 1985) and coincident with the final disappearance of the larval adipocytes, suggesting a transfer of their fat content to the adult fat body. Starvation and refeeding also decreased and increased cell size, respectively. It is likely, therefore, that adult fat body remodeling can take place only through changes in cell size, not in cell or nuclear number. However, further experiments should test the possibility that stimuli such as excess nutrition, damage or traumatic tissue loss might induce remodeling or regrowth through endoreplication, polyploid mitosis, depolyploidizing divisions or reactivation of cell fusion. Additionally, in light of the possibly low plasticity of the adult fat body, an interesting question to ask is whether the diet of the larva could imprint the metabolic status of the adult by affecting adult fat body development, for instance by influencing the initial number of precursors or their proliferative potential.

Besides characterizing adult fat body development, we importantly provide evidence of its physiological significance, for which conclusive prove had remained elusive. We found that flies where the adult fat body was missing or reduced (upon *srp* or *htl* knock down) displayed accumulation of neutral lipids in oenocytes and decreased viability upon starvation, demonstrating an essential role of the adult fat body as a lipid store and energy reserve. Consistent with this, starvation induced adipocyte autophagy and reduction in fat content, both reversible upon refeeding. By generating flies specifically lacking adult fat body, our study opens new avenues to systematically research adipocyte function in the adult, including roles beyond storage and metabolic regulation, for instance in endocrine signaling, matrix production, detoxification, immune responses, reproduction, and mating and feeding behaviors.

## ACKNOWLEDGMENTS

We thank Deborah Keiko-Hoshizaki for anti-Srp antibody. We also thank the Bloomington *Drosophila* Stock Center (BDSC), Vienna *Drosophila* RNAi Center (VDRC), Tsinghua Fly Center (THFC), Junhai Han, Yulong Li and Bing Zhou for providing fly strains, and José Martos-Marqués for technical advice on time-lapse imaging. This work was supported by grants 32150710524 and 91854207 from the Natural Science Foundation of China.

## AUTHOR CONTRIBUTIONS

All authors conducted research. Y.L., Y.H., K.Y., X.C. and J.C.P.-P. analyzed the data. J.C.P.-P. wrote the manuscript with input from all authors.

## METHODS

### *Drosophila* genetics

Standard fly husbandry and genetic methodologies were used to obtain the required genotypes for each experiment (see Table S1 for a detailed list of experimental genotypes). Fly strains and genetic crosses were maintained on standard medium prepared in our laboratory with yeast (24.5 g/L), cornmeal (50 g/L), agar (10 g/L), white granulated sugar (7.25 g/L), brown granulated sugar (30 g/L), propionic acid (4 mL/L), methyl-4-hydroxybenzoate (1.75 g/L) and absolute alcohol (17.5 mL/L). Pupae were staged by collection at the white pupa stage (0 h APF). Adults were staged by collection of newly eclosed animals from vials emptied at least 4 hours before (day 0 adult). Only males were imaged and analyzed, except for the survival experiment in Fig. 7D, in which both males and females were separately subjected to starvation. The GAL4-UAS system was employed to drive UAS transgene expression under the control of GAL4 drivers. Crosses were maintained at 25°C except for lineage tracing experiments involving *tub-GAL80*^*ts*^, in which cultures were maintained at 18°C until transferred to 30°C to initiate GAL4-driven gene expression. In lineage tracing experiments, expression of the yeast recombinase Flp driven by *twi-GAL4* or *Mef2-GAL4* excises an FRT-flanked sequence in a GAL4 flip-out cassette (Ito et al., 1997), turning on permanent, inheritable GAL4 expression in the affected cells and their progeny even after *twi-GAL4* or *Mef-GAL4* have ceased to be expressed in them. The time of labelling was additionally controlled with thermosensitive GAL4 repressor tub-GAL80^ts^ by staging and transferring animals from 18°C to 30°C at L1, L2, L3 or white pupa stage (0 h APF). The following strains were used:

*w*^*1118*^ (BDSC, 3506),

*w ; OK6-GAL4* (BDSC, 64199),

*w ; UAS-GFP*.*S65T* (BDSC, 1521),

*w ; UAS-myr-RFP / TM6B* (BDSC, 7119),

*y w ; UAS-mCD8-GFP* (BDSC, 5137),

*w ; UAS-GFP*.*nls* (BDSC, 4776),

*w ; UAS-mCD8-RFP* (BDSC, 32219),

*BM-40-SPARC-GAL4* (gift from Hugo Bellen),

*w twi-GAL4* (BDSC, 914),

*y w ; Mef2-GAL4* (BDSC, 27390),

*w ; Cg-GAL4* (BDSC, 7011),

*w ; ppl-GAL4* (gift from Pierre Leopold),

*y w ; act5c-y+-GAL4 UAS-GFP*.*S65T / CyO* (BDSC, 4411),

*y w ; UAS-Flp* (BDSC, 4539),

*w ; sco / CyO ; tub-GAL80*^*ts*^ (BDSC, 7018),

*y w ; UAS-mCherry.Atg8a* (BDSC, 37750),

*y v sc sev ; UAS-srp.RNAi*^*TRiP.HMS01298*^ (THFC, THU1529),

*w ; UAS-srp.RNAi*^*VDRC.v35578*^ (VDRC, v35578),

*y v sc sev ; UAS-srp.RNAi*^*TRiP.HMS01083*^ (BDSC, 34080),

*w ; UAS-htl.RNAi*^*VDRC.v6692*^ (VDRC, v6692),

*w ; UAS-htl.RNAi*^*VDRC.v27180*^ (VDRC, v27180),

*y w ; UAS-htl.DN.M ; UAS-htl.DN.M* (BDSC, 5366),

*w ; UAS-htl.lambda.M* (BDSC, 5367),

*y v ; UAS-pyr.RNAi*^*TRiP.HMJ30113*^ */ CyO* (BDSC, 63547),

*y w ; UAS-ths.RNAi*^*VDRC.v102441*^ (VDRC, v102441),

*w ; UAS-ths.RNAi*^*VDRC.v24536*^ */ TM3* (VDRC, v24536),

*w ; GMR93H07-GAL4* (BDSC, 40669),

*w ; ths*^*MI07139-TG4.1*^ */ CyO ; MKRS / TM6B* (BDSC, 77475),

*w ; elav-GAL80* (gift from Bing Zhou),

*y w ; act5C-GAL4 / TM6B* (BDSC, 3954) and

*w ; UAS-GFPS65C.αTub84B / CyO* (BDSC, 7374).

### Tissue dissections

To dissect pupal abdomens, we attached pupae to glass slides through their ventral sides using double-sided sticky tape. Then, we proceeded to open and peel the pupal case with fine tip forceps, pull out the animal carefully and transfer it to a Sylgard plate filled with PBS for dissection. Using dissection scissors, we separated the abdomen from the thorax and cut open the abdomen on its ventral side. Afterwards, we removed guts, gonads, larval fat body and other inner tissues using forceps. Abdomens were then fixed in 4% PFA for 15 min and washed twice in PBS for 15 min. After this, abdomens were either processed for tissue staining (see below) or mounted flat for direct observation with their inside surface up on a glass slide in a drop of DAPI-Vectashield (Vector Laboratories). A similar strategy was used for dissecting adult abdomens, only flies were immobilized by anesthetization with CO_2_, not stuck to a glass slide wit tape.

### Tissue stainings

For neutral lipid stainings, fixed pupal and adult abdomens were stained in BODIPY 493/503 (1:3,000 dilution of a 1 mg/mL stock, Life Technologies) in PBS for 1 h at room temperature and washed twice in PBS for 10 min before mounting in DAPI-Vectashield. For double staining of neutral lipids and actin cortex (Fig. 6B), we stained fixed pupal and adult abdomens with Texas-Red phalloidin (1:100, Life Technologies) and BODIPY 493/503 (1:3,000 dilution of a 1 mg/mL stock, Life Technologies) in PBT (PBS containing 0.1% Triton X) for 2 h at room temperature, followed by PBS washes (3 × 20 min) before mounting in DAPI-Vectashield. For anti-Srp antibody staining (Fig. 2A), fixed samples were blocked in PBT-BSA (PBS containing 0.2% Triton X-100 detergent, 1% BSA, and 250 mM NaCl) for 1 h, incubated overnight with anti-Srp primary antibody (1:200) in PBT-BSA at 4 °C, washed in PBT-BSA (3 × 20 min), incubated for 2 h in anti-rabbit IgG conjugated to Alexa-555 (1:200, Life Technologies) in PBT-BSA at room temperature, and washed in PBS (3 × 10 min). Samples were finally mounted in DAPI-Vectashield.

### Imaging of fixed tissues and analysis

Images of fixed adult abdomens stained with BODIPY in Fig. 2B and Fig. 4B-E, G and H were acquired in a Zeiss Axio Imager D.2 epifluorescence microscope using a 10 x / NA 0.3 objective. Other images of fixed pupal and adult abdomens were acquired with a Zeiss LSM780 upright confocal microscope using 10 x / NA 0.3, 20 x / NA 0.8, 40 x / NA 1.2 (water) or 63 x / NA 1.4 (oil) objectives. Nuclear counts in Fig. 6C were performed using the Multi-point tool in ImageJ-FIJI software. These counts were conducted on confocal stacks of images showing nuclear DAPI signal and plasma membrane *OK6>mCD8-GFP* (pupae) or phalloidin (adults). Three individuals were analyzed for each developmental time point. For nuclear ploidy estimation in Fig. 6E, confocal stacks of DAPI images were outlined and labelled with the Surface function in Imaris 9.8.1 software (Bitplane) and total DAPI signal inside the nucleus was computed. Ploidy was calculated with reference to the average DAPI fluorescence value of diploid (2n, 2C) blood cells.

### Live imaging and analysis

For live imaging of adult fat body formation during metamorphosis, pupae were removed completely from the pupal case with forceps and deposited on a glass-bottomed dish with a small drop of halocarbon oil 700 (Sigma) placed between the glass and the area to image. At the time of imaging, the glass-bottomed plate was inverted, leaving the animal hanging from the glass, attached to it by the surface tension of the oil. To maintain humidity, a piece of paper tissue soaked with water was located inside the dish. Imaging was conducted at a room temperature of 23°C in an upright Zeiss LSM780 confocal microscope using a 10 x / NA 0.3 objective for recordings of precursor migration (Video S1-S5), counting of *OK6>srp*^*i*^ precursors (Fig. 2D) and documentation of *htl* and *ths* expression (Fig. 4F and I). Number of precursors at 30 h APF and 36 h APF in wild type and *OK6>srp*^*i*^ animals were counted using the Multi-point tool in ImageJ-FIJI software. A 40 x / NA 0.95 (air) objective was used for recordings of precursor fusion (Video S6). From confocal stacks, maximum intensity projections or standard deviation projections were obtained using Zeiss Zen software. In recordings of precursor migration, 60 to 65 confocal sections were acquired per time point with a z-step of 1.7 to 2.0 μm at intervals of 4 min. In ventral view recordings and imaging in vivo (Fig. 1D, Fig. 2C, Fig. 3A and Fig. 4F), the legs were carefully displaced anteriorly, out of the imaging frame. For recordings of precursor fusion, 16 confocal sections were acquired per time point with a z-step of 1.2 μm at intervals of 5 min. Videos of maximum intensity or standard deviation projections were imported into Imaris 9.8 software to analyze precursor migration. The Spots tool was used to track cells and generate trajectories with the following parameters: 20 μm maximum distance between successive time points, 2 time points maximum gap size and 10 time points minimum track duration. Raw data for cell position was exported from Imaris into Excel, where we obtained displacement angles and speed through basic trigonometric calculations.

### Starvation assays

Starvation/refeeding experiments in Fig. 7B and C were conducted by placing 2 day male adults on starvation medium (2% agar medium) for 3 days before dissection. For refeeding, animals starved as above were transferred back to standard medium and then dissected. For survival tests in Fig. 7D, 2 day adults were placed on starvation medium and the number of surviving flies were counted at 12 h intervals. Three replicates of the experiment were performed separately for males and females, each replicate consisting of two vials containing 15-20 animals. Significance of differences with the control was tested through Mantel-Cox tests using GraphPad Prism 8. All differences were significant (****; p< 0.0001).

## SUPPLEMENTAL MATERIAL

**Video S1. Migration of adult fat body precursors into the abdomen**

**Video S2. Spreading of adult fat body precursors**

**Video S3. Expression of *htl-GAL4* in adult fat body precursors**

**Video S4. Spreading of *htl*^*DN*^ and *htl*^*CA*^ adult fat body precursors**

**Video S5. Detachment of *htl*^*DN*^ adult fat body precursors**

**Video S6. Fusion of adult fat body precursors**

**Table S1. Detailed genotypes**

